# Red light corrects neonicotinoid induced immunodeficiency and impaired respiration in poisoned bumblebees

**DOI:** 10.1101/2020.04.27.063925

**Authors:** Michael B Powner, Graham Priestley, Chris Hogg, Glen Jeffery

**Affiliations:** Centre for Applied Vision Research, City, University of London, UK; Institute of Ophthalmology, University College London, UK; Clinical Forums Ltd, Chichester, UK

**Keywords:** Photobiomodulation, Imidacloprid, Insect, Immunocompetence, ATP

## Abstract

Neonicotinoid pesticides undermine mitochondrial function in insects including bumblebees, reducing ATP, mobility and leading to death. They also reduce bumblebee immunocompetency leaving them vulnerable to pathogen attack. This undermines key pollinators critical in the agricultural economy. However, 670nm light exposure improves mitochondrial function undermined by age or disease, increasing respiratory chain efficiency, improving ATP production, mobility and survival in bumblebees and fruit flies. Here we show that 670nm restores immunocompetence, improving hemocyte counts and hemolymph anti-microbial action. Additionally, we measure whole body respiration *in vivo* in individual bumblebees revealing that it is a functional metric of both neonicotinoid impact and light induced mitochondrial protection. Critically we show that only 1 min 670nm exposure is sufficient to correct respiratory deficits induced by pesticide and restore normal immune ability. Longer exposures are not more effective. Further, single 1 min exposure protects respiration and immunity for approximately 3-6 days. Hence, 670nm impact is not dose dependent but switch like.

These data provide a compelling rational for 670nm application to protect pollinators on which a major part of the agro economy is based and who are being challenged by aggressive pesticide application.

## Introduction

Bee populations are in decline worldwide (Watanabe, 1994; Kosior et al., 2017; Nieto et al., 2014). This has a significant effect on ecosystem stability and pollination (Potts et al., 2010; Breeze et al., 2011; Ollerton et al., 2011; Hung et al., 2018; Fontaine et al., 2006; Klein et al., 2007). Factors suspected to contribute to this decline include, parasites and pathogens (Genersch et al., 2010) and pesticide exposure (Sandrock et al., 2014; Goulson et al., 2015). Neonicotinoid insecticides, are key contributors to reduced bee survival. They undermine mitochondrial function by overstimulating acetylcholine receptors (Moffat et al. 2015; Moffat et al,. 2016). The consequence is bumblebees have reduced ATP production, impaired visual function and restricted mobility resulting in reduced ability to feed leading to death (Lambin et al., 2001; Nauen et al., 2001; Suchail et al., 2001; Medrzycki et al., 2003; Colin et al., 2004; Powner et al., 2016). Neonicotinoids also suppress bee immunity rendering them vulnerable to pathogens (Brandt et al., 2016; Di Prisco et al., 2013),

Mitochondria are key players in ageing, disease and immunity. However, when mitochondrial function is undermined by ageing or disease it can be improved by long wavelength light that mitochondria absorb. In diverse species this improves their membrane potential, increases the efficiency of the electron transport chain and corrects compromised ATP production (Begum et al., 2013; Kokkinopoulous et al., 2013; Gkotsi et al., 2014). In insects it also improves retinal function, mobility, cognitive function including memory and extends average lifespan (Powner et al., 2016; Weinrich et al., 2017; Begum et al., 2015). Hence, in both bumblebees and flies daily exposure to 670nm light reduces death rates.

670nm protects bumblebees from Imidacloprid, a major neonicotinoid pesticide, when given for 15 mins twice daily. This corrects damaged retinal function, mobility and survival (Powner et al., 2016), but if this is to be used effectively, a greater understanding of the overall impact of 670nm light is needed in terms of its influence over immune-competence. Further, there is a need to understand the required metrics of exposure and their influence over time in bumblebees damaged by pesticide. Hence, here we examine for the first-time individual bumblebee respiration in response to insecticide poisoning and its protection by 670nm light over time and changes in damaged immune competence and its potential correction. Also, using similar metrics we ask how long improving single exposures of 670nm last. These data reveal significant improvement in the bumblebee’s immune response and respiration with brief light exposure providing a platform for treatment in the field.

## Materials and Methods

### Animals

Bumblebees (Bombus terrestris) were obtained in commercial colonies from Koppert UK. Experiments were undertaken in summer months. Bumblebees were maintained *ad libitum* on 50% sucrose solution in water and pollen.

### Exposure to Imidacloprid and/or 670nm light

Bumblebees were transferred from colonies and placed in 3L transparent plastic containers under standard 12/12 light dark cycles. Those exposed to Imidacloprid were given 10nM Imidacloprid in 50% sucrose solution in water. 670nm was delivered by specific 670nm light emitting LEDs. The spectral composition and energy output of these were checked before and after use. The energy levels in light exposures were a total of 40mw/cm^2^ from two light sources at either end of the container.

Intervention comparisons experiments had 4 groups; control, Imidacloprid, Imidacloprid + 670nm light and 670nm light on its own, identical to Powner et al. (2016). Here, 670nm light exposure was given twice daily by illumination from above with each exposure being 15 mins, and 12 h between exposures. Those treated with Imidacloprid were exposed continuously throughout via the sugar water.

To determine how long 670nm light exposure was needed to induce a positive biological response and once withdrawn how long its positive impact lasted, a time series of experiments were undertaken. Here bumblebees were again transferred from colonies and placed in 3L transparent plastic containers under standard 12/12 light dark cycles. Those exposed to Imidacloprid were given 10nM Imidacloprid in 50% sucrose solution. After Imidacloprid exposure (or control) single bumblebees were isolated into clear 12ml plastic test tubes, sealed in the tube using a gas permeable plastic bung. 670nm LED devices were placed either side of the test tube to deliver the required 670nm duration, at 40mW/cm^2^, which did not increase local temperature. Bumblebees had space to move up/down within the field of light. Bumblebees were returned to their 3L containers and respiration or immunocompetency experiment protocols followed.

### Measuring individual bumblebee respiration

This was undertaken to determine the impact of Imidacloprid on whole body respiration and how this was altered by 670nm light or immune challenge.

#### Respiration rate measurements

Whole body respiration was measured using a protocol adapted from Yatsenko et al. (2014). In brief, bumblebees were individually transferred into 12ml plastic test tubes. They were sealed in the tube using a gas permeable plastic bung. Soda lime (Sigma-Aldrich, Dorset, UK) was placed within the tube and the tube sealed with an airtight rubber bung. Sealed tubes had a syringe needle (19 gauge) forced through the base connected to a length of 1mm diameter clear tubing, the other end of which was placed in an ink bottle. This was identical to the procedure used by Weinrich et al (2018. See their Fig 2). In this sealed environment, when bumblebees expired CO_2_ it was absorbed by the soda lime generating a drop in internal pressure that could be measured by the volume of ink passing up the tube. Respiration was measured this way over 1h, and the average respiration rate per minute calculated.

To determine how long a period of 670nm light exposure was needed to improve respiration, and once withdrawn, how long its positive impact lasted, a time series of experiments were undertaken. Progressive exposures of increasing duration of 670nm were given at 0, 0.5, 1, 5, 15 and 60mins and respiration monitored in real time post 670nm exposure. To determine how long the positive impact of the 670nm light exposure remained effective, bumblebees were exposed to the minimal exposure period that was significantly improved respiration and then returned to the 3L containers and removed for respiration experiments at progressive periods at 0, 24, 48, 96, 144, 168 and 192h.

Bumblebee respiration rate was also measured after simulation of microbial insult. Bumblebees were maintained with/without exposure to Imidacloprid or 670nm as detailed previously for 4 days. The immune system of each bumblebee was then challenged by the injection into the abdomen of 1 μl of heat-inactivated *Escherichia coli* (OD 0.5). After a 24h period of immune response the respiration rates were measured, on day 5, as detailed previously.

### Determining bumblebee immune system health

The following were undertaken to determine the impact of Imidacloprid on immunocompetency and how this was altered by 670nm light.

#### Haemolymph collection

Haemolymph was collected using the protocol of Borsuk at al., (2017). Briefly; heads of bumblebees were swabbed with 70% ethanol and left to evaporate. An antenna of the bumblebee was detached and haemolymph outflow induced by pressing the abdomen. A bead (~5 µl) of haemolymph formed at the antenna base and collected with a pipette. This was immediately transferred to an Eppendorf and kept in ice to prevent melanisation. For haemocyte counts, the haemolymph was used immediately. Haemolymph destined for inhibition zone assays was stored at −80°C for 1 week until the assay was performed.

#### Total haemocyte count

1 µl of hemolymph was transferred to an Eppedorf tube containing 4 µl phosphate buffered saline and stored on ice. Diluted haemolymph was transferred to a hemocytometer (Burker chamber); haemocytes were counted (an average of three chambers per bumblebee were recorded) under an inverted phase contrast/fluorescent microscope. Hoechst (33258, Sigma-Aldrich, Dorset, UK) staining was used to confirm nucleus presence and cellular identity.

#### Inhibition-zone assay

Following 1 or 4 day exposure to Imidacloprid and/or/neither 670nm, the immune system of the bumblebee was challenged by the injection into the abdomen of 1 μl of heat-inactivated *Escherichia coli* (OD 0.5). After 24h hemolymph was collected as previously described (now on day 2 and 5 total exposure to varied conditions, respectively). Agar plates (9 cm diameter) containing LB medium (L3147, Sigma-Aldrich, Dorset, UK) were spread with 0.2 ml of fresh overnight cultures of *Micrococcus flavus* bacteria (OD 0.5). 1.5 μl aliquots of haemolymph were pipetted onto sterile 3 mm discs of blotting paper on the surface of the agar plate. The plates were incubated at 37°C for 24h and the diameter of each inhibition zone was measured using digital Vernier calipers. Data from each bumblebee was the average of two replicates.

### Statistical Analysis

Kruskal-Wallis H test and Mann Whitney U test were used to assess significance between groups. Error bars are Standard Error of the Mean (SEM).

## Results

### Immunocompetence

Haemocytes play a key role in the invertebrate immune system and total haemocyte counts are a metric of insect cellular immunocompetence (Wilson et al., 2002; Wilson-Rich et al., 2008). We quantify hemocyte number and their ability to inhibit bacterial activity when plated down *in vitro* after 2 and 5 days Imidacloprid exposure. The rational for these time periods is that 2 days exposures are similar to current literature (Brandt et al., 2016; Collison et al., 2018) and 5 days exposures correlate with the time period used in our previous study (Powner et al., 2016) and by others (Collison et al., 2018). In both cases this Imidacloprid exposure significantly reduced haemocyte numbers, halving their number compared to controls. However, twice daily, 15 minute exposure to 670nm light corrected this deficit. Exposure to 670nm light alone had no impact on hemocyte number. Extending exposure of Imidacloprid to 5 days did not change this result (Figure 1).

**Figure 1.**
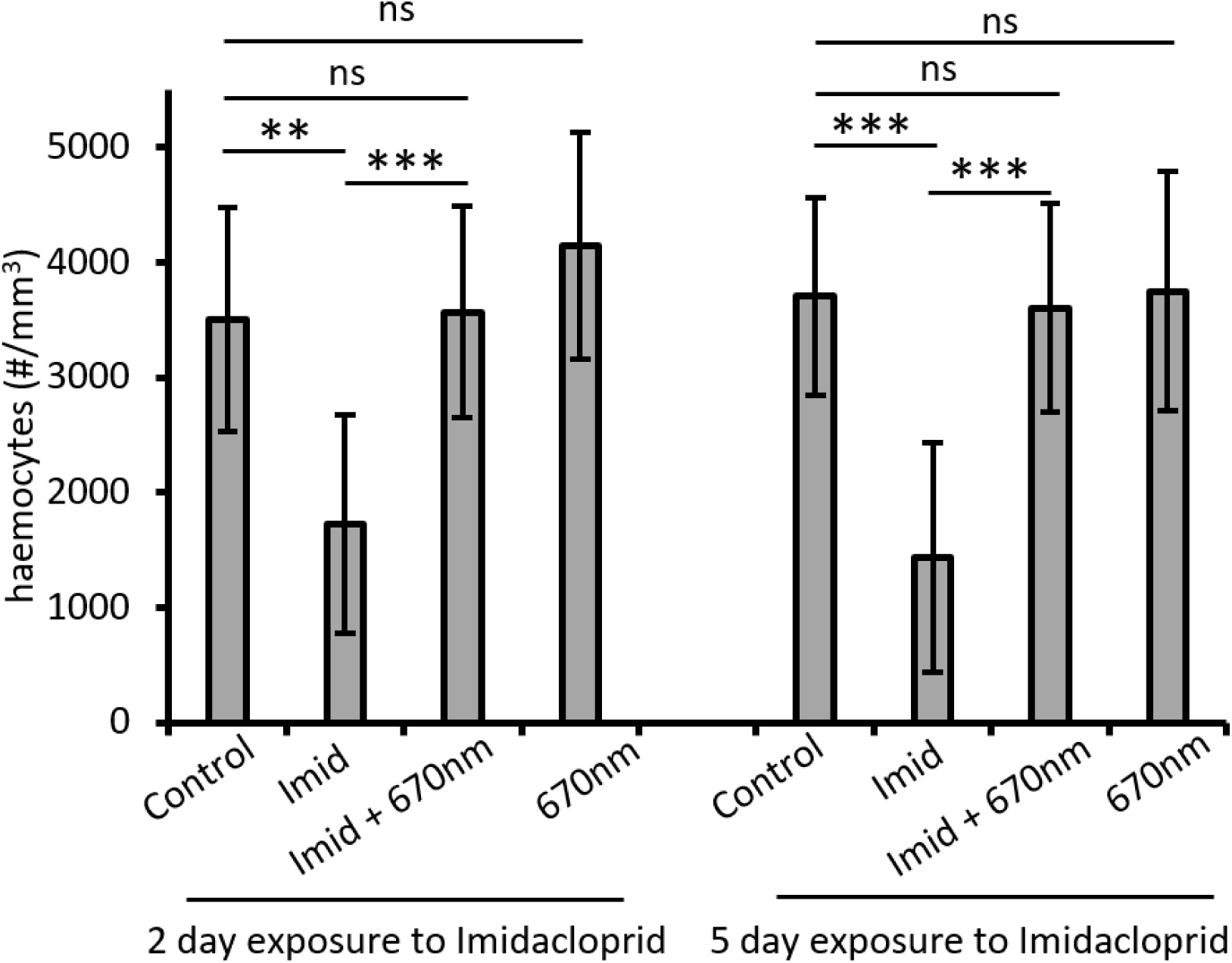
Haemocyte counts were used as a metric of immune health. Bumblebees were either exposed to Imidacloprid and or 670nm light or compared against controls. Exposures were for either 2 or 5 days and counts were made 2 days later. In both cases Imidacloprid had a detrimental effect on hemocyte numbers. This decrease was fully corrected with twice daily exposure to 670nm light. Abbreviations. ***; p < 0.005, ns; no significance, Imid; Imidacloprid. Bumblebees, ≥ 20 bees per group.

Immune competence was further investigated after bacterial challenge. Anti-microbial activity of haemolymph was tested *in vitro* 24h after animals were injected with dead bacteria. Here the size of the inhibitory zone induced by the haemolymph was measured similar to Brandt et al. (2016) as a measure. The antimicrobial activity of bumblebee haemolymph was significantly reduced in Imidacloprid exposed bumblebees following 2 and 5 day exposure to the insecticide. The size of the inhibition zone size decreased by approximately 35% after 2 day exposure to Imidacloprid compared to controls. This decrease in immunocompetence was similar following 5 day exposure to pesticide, indicating that the immune system is maximally compromised after 2 days. The reduction in immunocompetence induced by Imidacloprid was completely corrected by daily 670nm exposure at both 2 and 5 days. 670nm light had no effect on the antimicrobial activity of haemolymph in control bumblebees over either time period. Hence, both in terms of haemocyte number and their immune competence, Imidacloprid exposure had a significant negative impact, but this was completely corrected by 670nm exposure (Figure 2).

**Figure 2.**
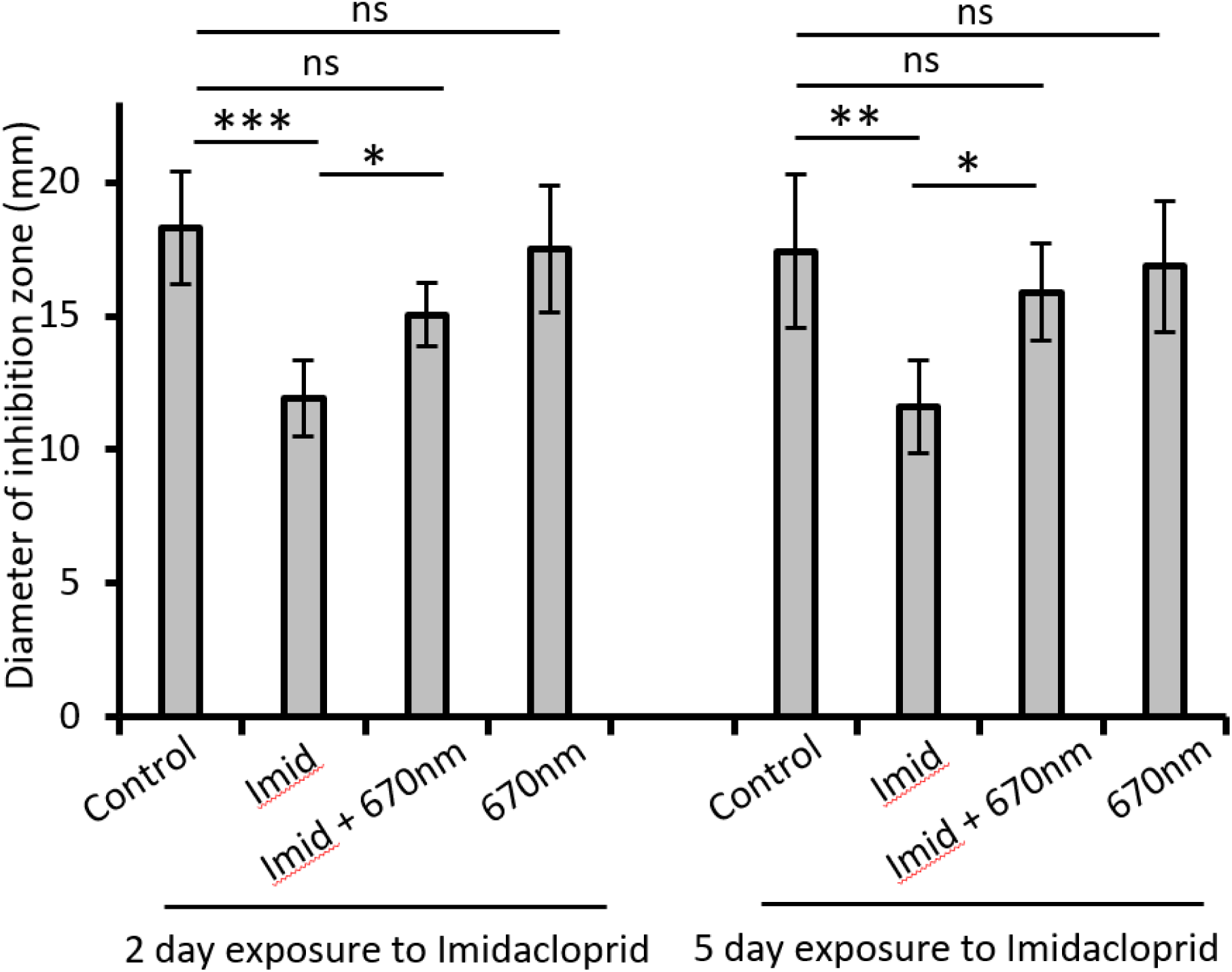
Antimicrobial properties of the circulating haemolymph was used as a metric of immunocompetence. After immune system challenge by *E.coli*, the haemolymph inhibited the growth of gram positive bacteria on agar plates after 2 or 5 days and counts undertaken 2 days later. The bumblebees immunocompetence of haemolymph is significantly reduced after exposure to Imidacloprid. This immune deficiency is partially restored with 670nm light exposure after 2 days exposure to both Imidacloprid and 670nm. Abbreviations **; p < 0.001, *; p < 0.05, ns; no significance, Imid; Imidacloprid. Bumblebees, ≥ 17 bees per group.

To determine the duration of 670nm exposure needed to restore compromised immunity and how long it remained effective a longitudinal series of tests were undertaken over different periods in bumblebees exposed to imidacloprid for 2 days. Exposure for 0.5min had no significant impact on haemocyte counts. However, with 1 min 670nm exposure there was a significant increase in number when counted 24h later. Longer exposures at 5, 15 and 60 mins also elevated haemocyte counts, but to levels no greater than found at 1 min. To reveal the duration of the effect on protecting hemocyte numbers following Imidacloprid exposure, bumblebees were then exposed to 1 min of 670nm light and their number counted over progressive hours. Significant improvements were present at 24 and 48h over controls, but their number declined at later stages and by 144h they were no different to controls. Hence 1 min of 670nm light is sufficient to protect haemocyte number following Imidacloprid exposure and this remains effective for 48h (Figure 3). 670nm exposure had no impact on haemocyte numbers in animals not challenged by Imidacloprid (data not shown).

**Figure 3.**
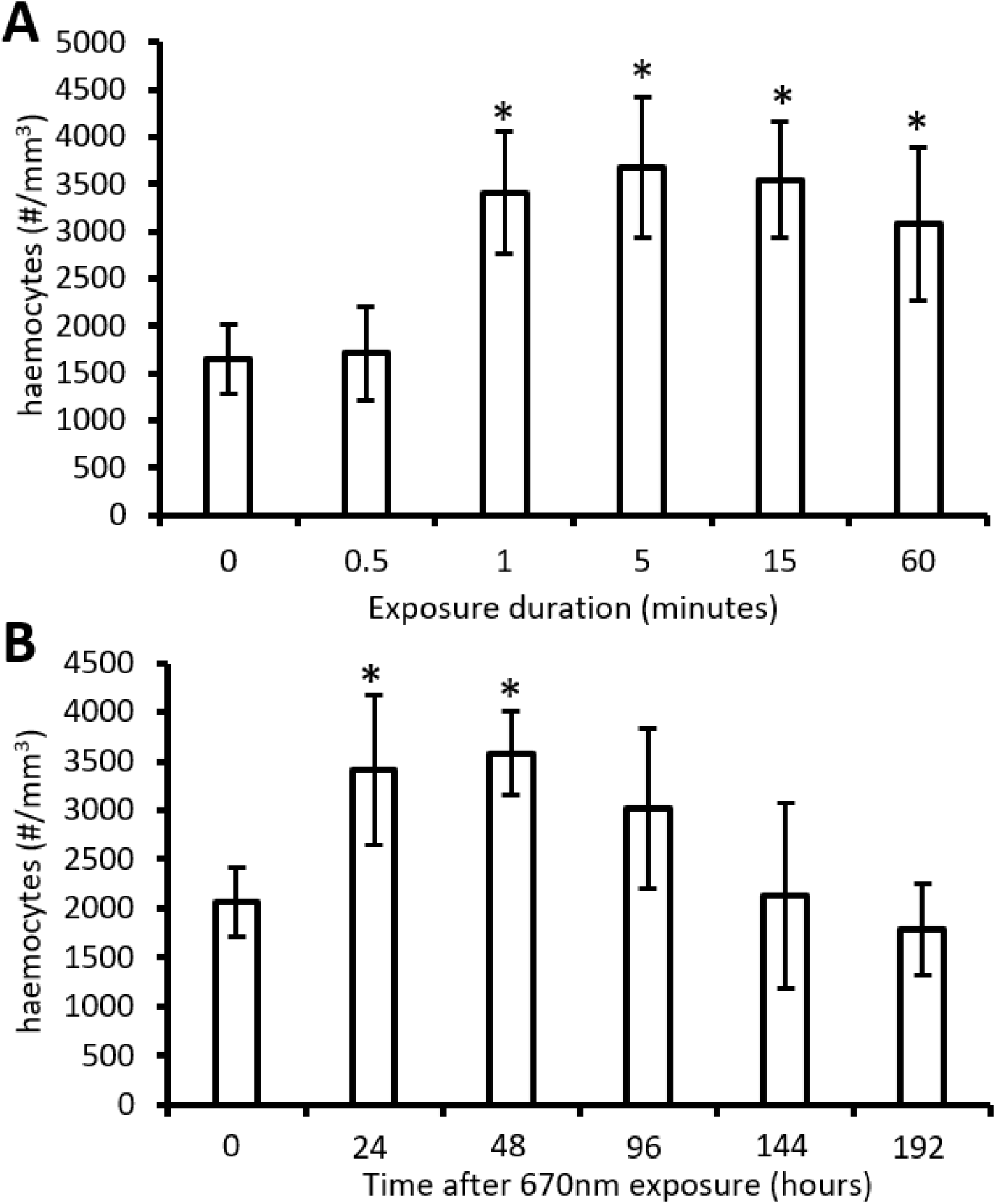
Exposure to 670nm restores haemocyte count of Imidacloprid treated bees, but the amount of 670nm needed and the duration of its impact are unknown. A. Bumblebees (≥ 20 bees per group) maintained on Imidacloprid were exposed to 670nm light for 0.5, 1, 5, 15 or 60mins with heamocytes counts carried out 24h later. Controls (0) were not exposed. Exposure for 0.5 min had no impact. But 1 min light exposure significantly increased haemocyte numbers. Longer exposures had similar effects. Bumblebees (≥ 22 bees per group) were exposed to a single 670nm for 1 mins (B) and haemocytes counted at 24h intervals after. Using bumblebees exposed to Imidacloprid; this single 1 min exposure restored haemocyte counts over for 24h and 48h, however a decrease was observed from 96h post exposure, with no significance difference from the control since afterwards. Abbreviations: *; p < 0.05.

### Respiration

Imidacloprid undermines mitochondrial function and hence likely impacts on respiration. We quantify whole body respiration in individual bumblebees by measuring their CO_2_ production. Here Imidacloprid had a significant impact reducing respiration. This reduction was corrected in bees exposed to 670nm light. Here the light induced a large increase in respiration above that found in controls. However, as this level of exposure was similar to that in our previous study that resulted in protected mobility and extended lifespan in insecticide treated bumblebees, the elevated respiration is likely to be protective (Figure 4)

**Figure 4.**
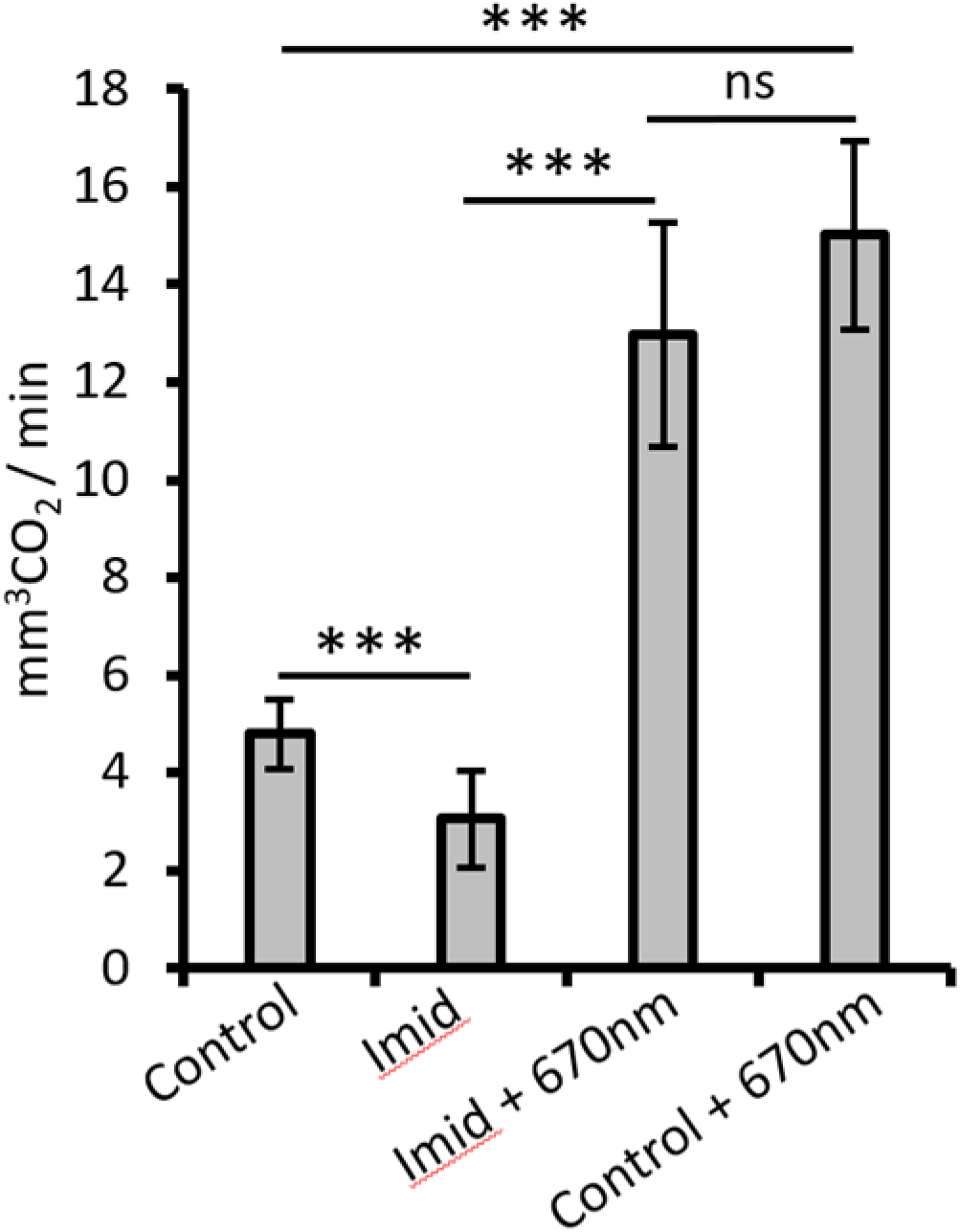
Bumblebee respiration was measured. They were either exposed to Imidacloprid and/or 670nm light or compared against controls. Imidacloprid and or 670nm exposure was for 5 days (≥ 18 bees per group). (A) In both types of bee Imidacloprid had a significant impact reducing respiration, but this was corrected by exposure to 670nm light. Abbreviations: ***; p < 0.005, ns; no significance. Imid; Imidacloprid

In an immune response, cellular energy demand increases. Hence, we ask if Imidacloprid undermines the ability of the animal to produce an appropriate immune response in terms of their respiration. Bumblebees were challenged with dead bacteria and 24h after their respiration rates were measured. Figure 5 shows that respiration in normal bumblebees is elevated by exposure to dead bacteria when compared to controls shown in Figure 4. However, this was significantly suppressed by Imidacloprid. Application of 670nm light in Imidacloprid treated bumblebees corrected the deficit. However, it was not increased in bumblebees injected with dead bacteria and also exposed to the light (Figure 5). This may be because the respiration rate in the control animals was close to maximal in this situation.

**Figure 5:**
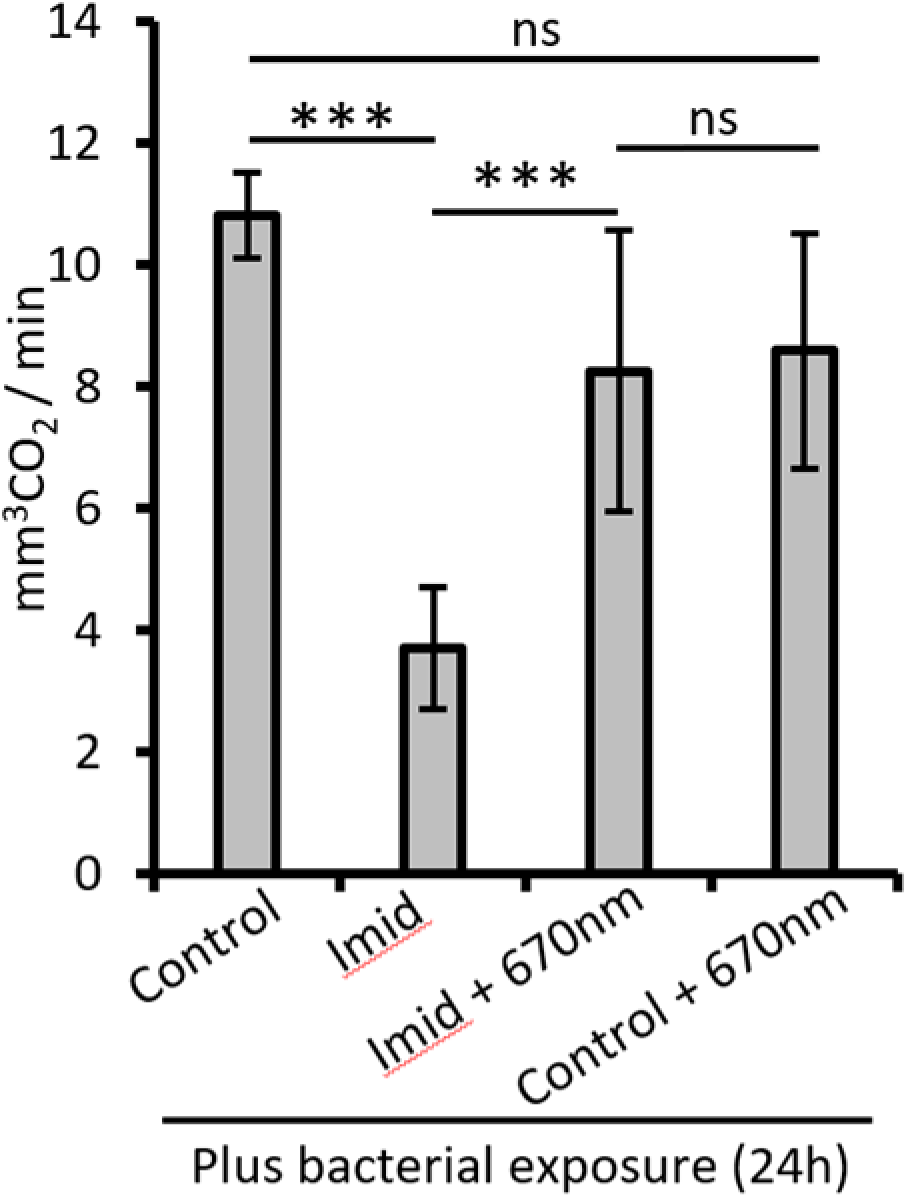
During the immune response cellular energy demands increase. Bumblebees were injected with dead bacteria and 24h later their respiration was measured. Controls who were injected but not exposed to any other manipulation have increased respiration over untreated bees (see Fig 4.). However, exposure to Imidacloprid after microbial insult significantly reduced respiration, which was returned following 670nm exposure. (≥ 22 bees per group). Abbreviations: ***; p < 0.005, ns; no significance. Imid; Imidacloprid

Our results reveal that 670nm can be used to improve respiration, but as with immunity we ask how much 670nm is needed and how long do its effects last. In bumblebees not exposed to Imidacloprid 0.5 mins exposure had no impact on respiration, but 1min exposure increased it significantly and this was the same for longer exposures. Hence, only 1 min is needed to improve respiration. Having established this, we use 1 min exposure and then examine respiration in the same bumblebees over 192h. Respiration remained elevated for 144h. This paradigm was then repeated in bumblebees exposed to Imidacloprid with largely similar results. 1min exposure to 670nm was sufficient to significantly improve respiration and this remained effective for 96h following insecticide exposure (Figure 6).

**Figure 6.**
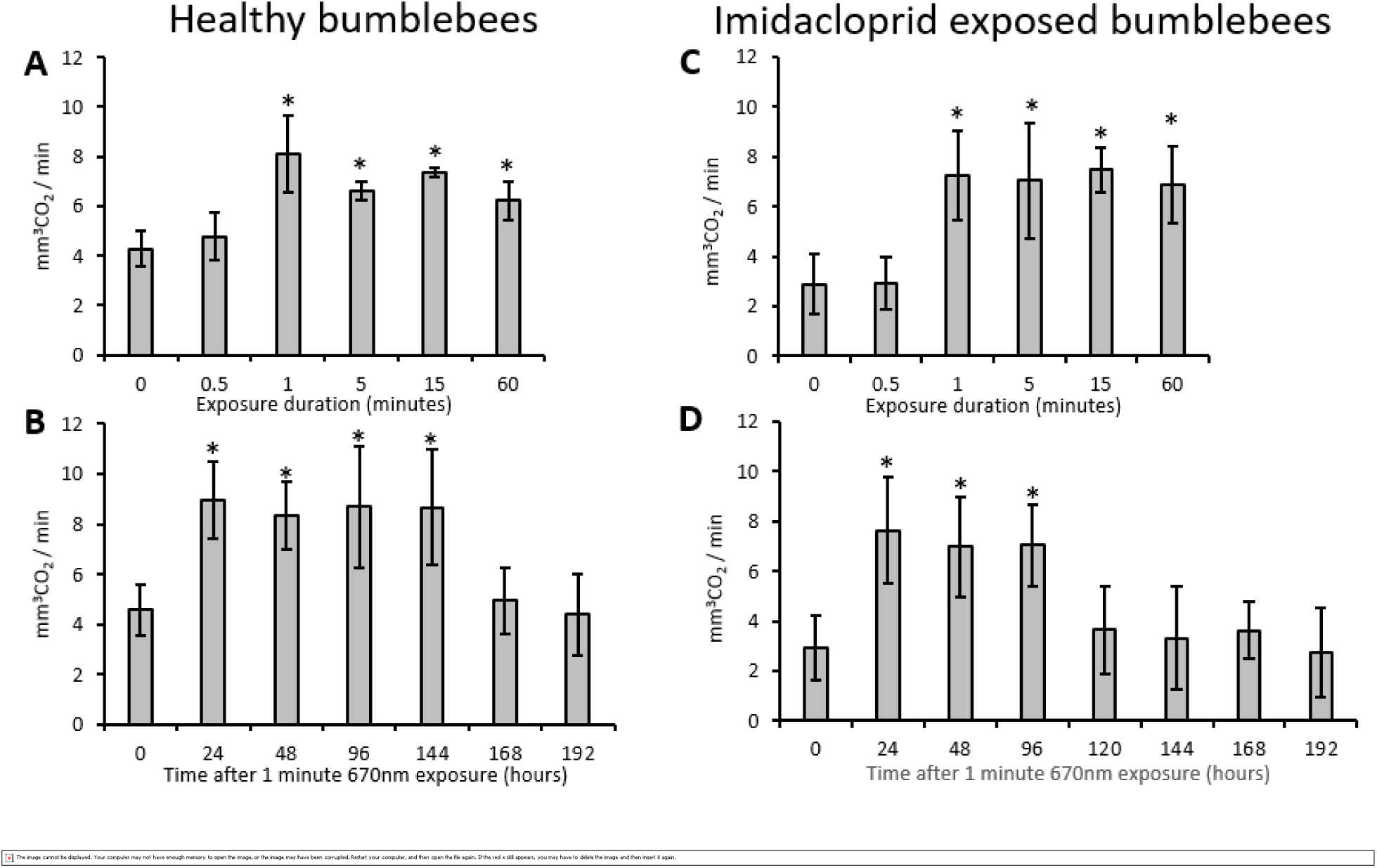
Exposure to 670nm increases respiration but the amount of light needed and the duration of the effect is unknown. A. Bumblebees (≥ 20 bees per group) were exposed to 670nm light for 0.5, 1, 5, 15 or 60mins and respiration measured. Controls (0) were not exposed. Exposure for 0.5 min had no impact on respiration. However, 1 min light exposure increased significantly respiration. Exposures for longer periods were similarly effective. Bees (≥ 18 bees per group) were exposed to a single 670nm for 1 mins (B) and respiration measured at 24h intervals after. This single exposure elevated respiration over a 144h period, but then declined rapidly afterwards. The amount of light needed and duration of effect were also determined in Imidacloprid exposed bees (C and D). Exposure of 0.5min had no impact of respiration, however durations between 1 and 60mins induced significant increase in respiration rate (C). A single 1min exposure of 670nm elevated respiration rate for 96h, a slight reduction in duration compared to the effect of 670nm on non-pesticide exposed bees. Abbreviations: *; p < 0.05.

## Discussion

This study shows that Imidacloprid, a neonicotinoid insecticide, directly undermines whole bumblebee respiration and immunocompetence. Further, that in both cases only brief exposure to 670nm light of approximately 1min restores both features and that the positive impact of such brief exposure lasts for 4-6 days. This strongly implies that each of these features has a common underlying mechanism that most likely resides with mitochondrial integrity. These metrics have not been previously explored in bee populations. While there are an increasing number of pesticides available, it is likely that any targeting mitochondrial function will respond positively to 670nm treatment via similar mechanisms.

Previously we have shown that 670nm light in Imidacloprid poisoned bumblebees restore damaged mitochondrial production of ATP, animal mobility and improves lifespan. As there is an intimate and complex relationship between mitochondrial function and immunity (Arnoult et al., 2009; Weinberg et al., 2015; Mills et al 2017), it is not surprising that this study has been able to reveal a positive impact on the immune response following recovery of mitochondrial function.

In spite of the overall improvements shown here that are consistent with a wide range of studies using experimental pathology across species (Fitzgerald et al., 2013) there are marked gaps in understanding. We have counted total haemocye number, but these are not a heterogeneous population and it is unclear which type is responding to the light (Marringa et al., 2014). Likewise, the restoration of total haemocyte number does not indicate a restoration of the exact immune system profile as the control bumblebees. There are also likely to be complex interactions between the antimicrobial properties of the haemolymph and the number of haemocytes that remain to be explored.

Our data also reveal that whole body respiration in bumblebees is a relatively simple and effective metric for assessing both the impact of Imidacloprid and also for its potential resolution by exposure to 670nm light. These experiments reveal that only 1 min exposure is sufficient to resolve undermined respiration and here the effect can last 144h. This is consistent with data from Drosophila, where single exposures of 670nm remained effective for 100h (Weinrich et al., 2018). Our data also show that 670nm light exposure increased respiration in both treated and untreated bumblebees well above the level found in control and it may be argued that this may be problematic. However, these exposures are associated correcting the deficits found following Imidacloprid exposure and also in normal animals with a marked increase in average lifespan (Powner et al., 2016). Consequently, it is unlikely that they are detrimental.

Weinrich et al. (2017) did much to reveal the changes that take place with 670nm light in aged Drosophila, showing improved metabolism in terms of triglyceride, glucose and glycogen storage and that this is associated with increased robustness when animals are physiologically challenged by chill. Data from this study also showed improved aged memory along with mobility and retinal function following 670nm exposure. This matches the improved mobility and retinal function also found in bumblebees exposed to 670nm after insecticide exposures. (Powner et al., 2016). Further, The debilitating impact of Imidacloprid on bumblebee function and its restoration by 670nm light finds symmetry in many mammalian studies where this light has been employed to reduce the impact of induced CNS pathology, particularly when this is associated either directly or indirectly with mitochondrial insult (Fitzgerald at al., 2013; Darlot et al., 2016). However, particularly pertinent to the application of 670nm light in bees generally is the finding that old flies lose their stereotypic patterns of navigation in an open environment, but that this rapidly returns after a single 670nm light exposure (Weinrich et al., 2018). Hence, is it likely that using 670nm light may help bees maintain their navigational abilities when foraging. As we know that 670nm light also improves memory and retinal function (Powner et al., 2016; Weinrich et al., 2017), its application to bee hives/colonies may offer significant advantages.

A key feature of our data is the absence of any conventional dose dependent effect following 670nm exposure. Relatively long exposure to 670nm was not better than 1 min. This appears to be a feature of long wavelength light in mammalian models of induced pathology where variable light exposures have very similar impact (Fitzgerald et al., 2013). Likewise, the effect of the light did not decline gradually over days, but appeared to terminate over a relatively short time. These data indicate that the underlying mechanism does not act in a graded manner but is rather switch like.

Our data show that only 1 min of light exposure had significant effect against Imidacloprid. This is potentially important because it is possible to expose honey bees that live in hives to this brief light when they pass in and out of the hive. An additional advantage of 670nm light is that it is beyond the bee’s visual range and consequently the animals are not disturbed by it (De Ibarra et al., 2014).

We have used bumblebees in our experiments because colonies of these animals are commercially available, which is not the case for honey bees. Further, these colonies are smaller than honey bee hives and more suitable to the lab environment. But we have used honey bees in respiration limited studies and the data generated are very similar to that of the bumblebee. Consequently, there is a strong reason to believe that treating honey bees with 670nm will be effective. But critically our experiments are lab based. However, Beefutres in France have adopted the technology we have used in this study in real field experiments and hence may provide the translation needed in honey bees in a natural setting. Here the data being generated are consistent with that produced in the lab, reinforcing the idea that long wavelength light is likely to be of value in sustaining this key pollinator.

## Author Contributions

M. Powner, and G. Jeffery designed research; M. Powner and G. Jeffery performed research; G. Priestley and Chris Hogg contributed reagents and technical advice; M. Powner and G. Jeffery analyzed data; M. Powner, T. G. Priestley and G. Jeffery wrote the paper.

## Acknowledgements

This research was supported by the BBSRC BB/N000250/1.

